# Target-decoy false discovery rate estimation using Crema

**DOI:** 10.1101/2023.06.18.545038

**Authors:** Andy Lin, Donavan See, William E. Fondrie, Uri Keich, William Stafford Noble

## Abstract

Assigning statistical confidence estimates to discoveries produced by a tandem mass spectrometry proteomics experiment is critical to enabling principled interpretation of the results and to assess the cost/benefit ratio of experimental follow-up. The most common technique for computing such estimates is to use *target-decoy competition* (TDC), in which observed spectra are searched against a database of real (target) peptides and a database of shuffled or reversed (decoy) peptides. TDC procedures for estimating the false discovery rate (FDR) at a given score threshold have been developed for application at the level of spectra, peptides, or proteins. Although these techniques are relatively straightforward to implement, it is common in the literature to skip over the implementation details or even to make mistakes in how the TDC procedures are applied in practice. Here we present Crema, an open source Python tool that implements several TDC methods of spectrum-, peptide- and protein-level FDR estimation. Crema is compatible with a variety of existing database search tools and provides a straightforward way to obtain robust FDR estimates.

## 1 Introduction

The goal of a typical proteomics mass spectrometry experiment, whether it is carried out on samples from humans, model organisms, or environmental samples, is to produce biological insights that can inform future experiments. The purpose of assigning statistical confidence estimates to such results is to allow scientists to assess the likely utility of such follow-up experiments by quantifying the estimated rate of false positives among the **discoveries**^1^ produced by the experiment.

By far the most common approach for estimating the **false discovery rate** (FDR) among spectrum identifications or peptide or protein detections from a mass spectrometry experiment is the target-decoy competition (TDC) framework.^1^ This approach involves searching the observed spectra against a database comprised of real (target) peptides and an equal number of decoy peptides, where the decoys may be created by reversing or shuffling the amino acids of the targets. FDR control among the identified spectra (**peptide-spectrum matches**, PSMs) is attempted by competing the target PSMs against the decoy PSMs, retaining only the best-scoring PSM per spectrum. The FDR among the optimal PSMs above a specified score threshold (known as the “accepted PSMs”) can then be estimated by taking the ratio of the numbers of accepted decoy versus target PSMs (Section 2.1). This procedure provably controls the FDR—the expected rate of false discoveries among accepted PSMs.^2^ Similar competition-based procedures have also been developed for FDR control among detected peptides or proteins.^3,4^

When formulating and proving the underlying mathematical theory of TDC,^2^ He *et al*. cautioned against the use of TDC at the PSM level, a warning that was recently reinforced.^5^ At the same time, both papers argued that TDC does manage to control the peptide-level FDR—the expected rate of false discoveries among reported peptides. Interestingly, the core idea of TDC, namely, that incorrect discoveries are equally likely to come from a target or decoy match, also plays a central role in the widely used knockoff framework for general FDR control,^6^ which was introduced by Barber and Candés in 2015.

Unfortunately, although TDC procedures for FDR control are relatively straightforward to implement, it is not uncommon for mistakes to be made in their application. Perhaps the most common problem is simply omission of implementation details from the methods section of published papers. Methodologically, one challenge is ensuring that every peptide in the target database has a corresponding isobaric peptide in the decoy database. This property is required for valid FDR control, yet as has been pointed out repeatedly in the literature,^3,7–10^ several common methods for producing a decoy database do not maintain equal-sized target and decoy databases. Another potential source of error is that, to accurately control the FDR, a “pseudocount” of +1 must be added to the number of accepted decoy PSMs.^2,11^ This pseudocount has a relatively small impact on studies with thousands of discoveries but can become consequential for studies with few discoveries or that aim for very strict FDR control. Many additional types of mistakes can be introduced when FDR control is carried out in conjunction with a machine learning post-processor, such as PeptideProphet^12^ or Percolator,^13^ or in complex analysis strategies in which subsets of the discoveries are analyzed separately.^14^ A general rule of thumb is that information about which discoveries are targets and which are decoys must not “leak” into the analysis pipeline prior to FDR estimation, but detecting such leakages can sometimes be challenging. Finally, an unfortunately common mistake is to misinterpret FDR estimates by first controlling the FDR in one set of discoveries, selecting a subset of those discoveries for analysis or follow-up, and then erroneously associating the original FDR estimate with the subset.^5,15,16^

A number of existing methods provide the ability to estimate the FDR on a set of detections. The MS-GF+ search engine implements a TDC procedure internally.^17^ Several proteomics data analysis toolkits, such as Crux,^18^ Pyteomics,^19,20^ and OpenMS,^21^ provide commands for estimating the FDR at the PSM or peptide levels. Furthermore, post-processor programs such as Percolator^13^ and PeptideProphet^22^ also estimate FDR on a set of detections, after first applying machine learning methods to re-rank the discoveries produced by a search engine. Recently, Debrie *et al*. created a software tool to empirically evaluate whether the assumptions that TDC requires are followed,^23^ but the tool itself does not provide FDR estimates.

As a step toward ensuring more uniform and accurate FDR control procedures in mass spectrometry proteomics, we have created Crema, an open source Python tool that computes FDR estimates at the PSM, peptide and protein levels. Crema works directly with a variety of widely used database search tools that produce search results in tab delimited, pepXML, or mzTab formats. We provide a tutorial case study to illustrate how to use Crema, as well as examples of how to use Crema with Tide,^24^ MS-GF+,^17^ Comet,^25,26^ MSFragger,^27^ and MSAmanda.^28^ We also show empirical results on a variety of publicly available datasets.

## 2 Methods

### Box 1

Key definitions

- **Peptide-spectrum match** (PSM): Assignment of a peptide from the database to an observed spectrum.
- **Discovery**: One observation drawn from the data. This can be a PSM, a detected peptide, or a detected protein.
- **Accepted discovery**: A discovery whose associated score is above a specified threshold.
- **False discovery proportion** (FDP): The proportion of accepted discoveries that are false positives (type I errors).
- **False discovery rate** (FDR): The expected (i.e., average) FDP.

### 2.1 PSM-level FDR control

FDR control is commonly performed at the PSM level following a database search. However, it has been demonstrated that PSM-level FDR control is problematic because it can be liberally biased.^2,5^ This bias occurs because one of the assumptions that is required for valid FDR control—namely, that the incorrect PSMs are independent of one another—is violated in practice.^2^ We therefore strongly recommend avoiding PSM-level FDR and instead use peptide-level FDR control. Nonetheless, due to the ubiquity of PSM-level analysis, Crema provides a procedure that attempts to perform PSM-level FDR control.

#### 2.1.1 PSM-level TDC

The goal of TDC carried out at the PSM level is to estimate the false discovery rate among a collection of PSMs produced by a database search engine. We are given a set *S* of *n* spectra, and we assume that the spectra have already been searched against a target database *𝒯* and a decoy database *𝒟*. Critically, the target and decoy databases must be the same size and must exhibit the same distribution of masses, so that each target peptide has a corresponding isobaric decoy peptide. For each spectrum we retain the top-scoring PSM across the target and decoy database, breaking any ties randomly. We refer to the scores of the optimal PSMs that involve target peptides as 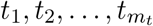, and of those that involve decoy peptides as 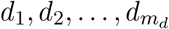, where *m*_*t*_ + *m*_*d*_ = *n*. We can then estimate the FDR among all PSMs that score greater than a specified score threshold *τ* (assuming larger scores are better) as

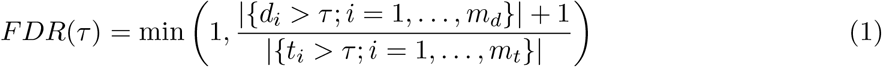

Intuitively, the denominator represents the number of discoveries of interest (the target PSMs), and the numerator is our decoy-based estimate of the number of false positives among those discoveries.

The formulation in Equation 1 differs from the one offered by Elias and Gygi^1^ in three ways. First, their formulation includes a factor of 2 in the numerator and includes both target and decoy PSMs in the denominator. This approach thus controls the FDR among the combined set of target and decoy PSMs. In practice, it typically is of more interest to control the FDR only among the targets, as in Equation 1. Second, the numerator in our formulation includes a +1 that is missing from the Elias and Gygi formulation. This +1 correction is analogous to +1 corrections in permutation testing, and it is required in order to achieve valid FDR control under the assumption that incorrect identifications are independently equally likely to be targets or decoys.^2,11,29^ In practice, this +1 correction will have a negligible effect except in the presence of very few discoveries or a very stringent FDR threshold. Finally, our formulation includes an enclosing min operation, which simply ensures that we do not report an FDR *>* 1.

### 2.2 Peptide-level FDR control

Aside from the aforementioned problematic nature of FDR control at the PSM level, FDR control at the PSM level, in practice, may not be as useful as controlling FDR at the level of peptides, especially if the primary conclusions are being drawn about peptides rather than spectra. Crema offers three previously described^5^ peptide-level FDR controlling procedures—psm-only, peptide-only, psm-peptide (previously called “PSM-and-peptide”)—all based on TDC. Empirical evidence suggests that psm-peptide offers the best statistical power, but Crema offers implementations of all three methods.

#### 2.2.1 PSM-only peptide TDC

The first method, PSM-only peptide TDC (“psm-only”), is both the traditional and most commonly used method for estimating peptide-level FDR. In this method, the direct competition is only done at PSM level: for each spectrum we compare its best matching decoy peptide with its best matching target peptide and we keep the higher scoring of the two PSMs. Note that ties are randomly broken and that this process is largely equivalent to searching each spectrum against the concatenated target-decoy database. Next, each target/decoy peptide is assigned the score associated with its best scoring PSM. Finally, with a slight abuse of notation, if we refer to the list of target peptides as 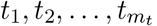, and similarly for decoys 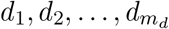 (in this case *m*_*t*_ = *m*_*d*_), then the peptide-level FDR can be estimated using Equation 1. This method can be used within Crema by setting “pep_fdr_type=psm-only” when calling the assign confidence function.

#### 2.2.2 Peptide-only peptide TDC

The second peptide-level FDR procedure, peptide-only, requires that each peptide in the target database is paired with its corresponding peptide in the decoy database. In this case the direct competition only takes place at the peptide level: each spectrum is separately searched against the target and the decoy databases to find its optimal target and decoy PSMs. Each target/decoy peptide is then assigned the score associated with its best scoring PSM, and from each target-decoy pair of peptides only the higher scoring peptide is retained. We note that ties are randomly broken. Referring to this filtered (winning) list of target peptides as 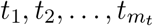, and similarly for decoys 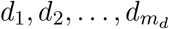, the peptide-level FDR is again estimated using Equation 1. We note this procedure is similar to the “picked protein” procedure described below. This method can be used in Crema by setting “pep_fdr_type=peptide-only” when calling the assign confidence function. The pairing between target and decoy peptides can be explicitly provided to Crema via the *pairing_file* parameter when reading the PSM files. This pairing file is a text file in which the first column contains a target peptide and the second column contains the corresponding paired decoy peptide. If Tide or Comet, within Crux, is used as the database search engine, the pairing file can be generated by setting “–peptide-list=T” when building the peptide index. Alternatively, for these search engines, the pairing file can be implicitly created, as described below. Otherwise, this file must be generated by the user.

In addition to an explicit pairing, a peptide pairing can be implicitly created if Crema is provided the search results of Tide or Comet. Comet generates decoy sequences by reversing a target peptide sequence, while keeping the first and last amino acid fixed. Therefore, the pairing between targets and decoys is self-evident. Tide can generate decoy sequences by either reversing or shuffling the internal amino acids of a peptide sequence, while keeping the first and last amino acid fixed. Crema infers the target-decoy matching based on amino acid composition, selecting matches at random as necessary.

Unfortunately, target and decoy peptides cannot be paired based on results from MSGF, MSAmanda, or MSFragger because these search engines create decoys by reversing or shuffling protein sequences. In order for a peptide pairing to be created, decoys must be generated at the peptide level since a paired target and decoy sequence must be isobaric to ensure they are always considered by the same set of spectra. To explain why protein reversal does not work, consider the toy protein sequence of “ACRMCK”. This protein will generate two peptides when digested by trypsin: “ACR” and “MCK.” If the protein is reversed, then it will generate three peptides: “K”, “CMR”, “CA.” None of these decoy sequences have the same mass as any of the targets, and therefore a pairing cannot be generated. Table 1 summarizes which types of FDR control can be carried out with which search engines.

**Table 1:** FDR control procedures implemented for each score function. Crema implements one, three, and one FDR control procedure at the psm, peptide, and protein level, respectively.

| score | PSM level | peptide level |  |  | protein level |
| --- | --- | --- | --- | --- | --- |
|  |  | psm-only | peptide-only | psm-peptide |  |
| XCorr | ✓ | ✓ | ✓ | ✓ | ✓ |
| XCorr p-value | ✓ | ✓ | ✓ | ✓ | ✓ |
| Tailor | ✓ | ✓ | ✓ | ✓ | ✓ |
| combined p-value | ✓ | ✓ | ✓ | ✓ | ✓ |
| Comet e-value | ✓ | ✓ | ✓ | ✓ | ✓ |
| MSGF+ | ✓ | ✓ |  |  | ✓ |
| MSFragger | ✓ | ✓ |  |  | ✓ |
| MSAmanda | ✓ | ✓ |  |  | ✓ |

#### 2.2.3 Peptide-level FDR with PSM competition

The third procedure, “psm-peptide” executes both PSM and peptide-level competitions. First, as in PSM-only, for each spectrum only the top matching target/decoy PSM is kept. Thereafter, it continues as in peptide-only: scores are aggregated to the peptide level by retaining only the top-scoring PSM per peptide, and a second round of competition is carried out, this time between paired target and decoy peptides. The FDR among the resulting detections is again estimated using Equation 1. Similar to peptide-only, because psm-peptide requires peptide-level pairing information, it is only compatible with Comet and Tide searches. This method can be used by setting “pep_fdr_type=psm-peptide” when calling the assign confidence function.

### 2.3 Protein-level FDR control

In addition to PSM- and peptide-level FDR control, Crema provides protein-level FDR control via the picked protein FDR procedure, which carries out competition at the protein level.^4^ Similar to the picked peptide approach, the picked protein procedure begins by performing PSM-level TDC to identify the top-scoring PSM per spectrum. Subsequently, peptides that occur in more than one protein in the database are removed. Using the unique peptides, each protein is assigned two scores: the maximum score (or minimum score, if a smaller score is better) associated with all of that protein’s target peptides, and the maximum score associated with all of that protein’s decoys. The final protein score is the maximum of these two values. Assuming that 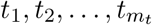 and 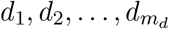 now refer to protein-level scores, the protein-level FDR is calculated using Equation 1.

### 2.4 Q-value calculation

For benchmarking purposes, Crema can also compute the q-value for each target detection *t*_*i*_, which can be a PSM, peptide, or protein. The q-value is defined as the minimum FDR threshold at which this detection would be accepted, i.e.,

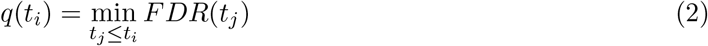

Note that we strongly discourage the use of q-values for anything other than benchmarking. In particular, selecting a desired q-value cutoff *β* by comparing the list of discoveries corresponding to different q-values does *not* control the FDR at level *β*. Practically, this means that the FDR threshold should be fixed prior to analysis.

### 2.5 Implementation

Crema is an open source project written in Python, and the source code is available on Github at https://github.com/Noble-Lab/crema with an Apache license. Crema can be installed using pip and run via a Python package API or through the command line. Installation instructions and usage documentation can be found at https://crema-ms.readthedocs.io. In addition to Python 3.6+, the following dependencies are required, and will be automatically installed via pip: numba, numpy, pandas, lxml, and pyteomics.

### 2.6 Dataset and database searches

For the experiment reported here, we used five LC-MS/MS runs, where each run was generated with a sample from a different species—castor plant, yeast, human, mouse, and *Escherichia coli*— from four different studies (Table 2). The raw files were downloaded from PRIDE^30^ and converted to mzML format using MSConvert version 3.0.^31^ The reference proteomes were downloaded from Uniprot^32,33^ (www.uniprot.org) in October 2021 and January 2022.

**Table 2:**
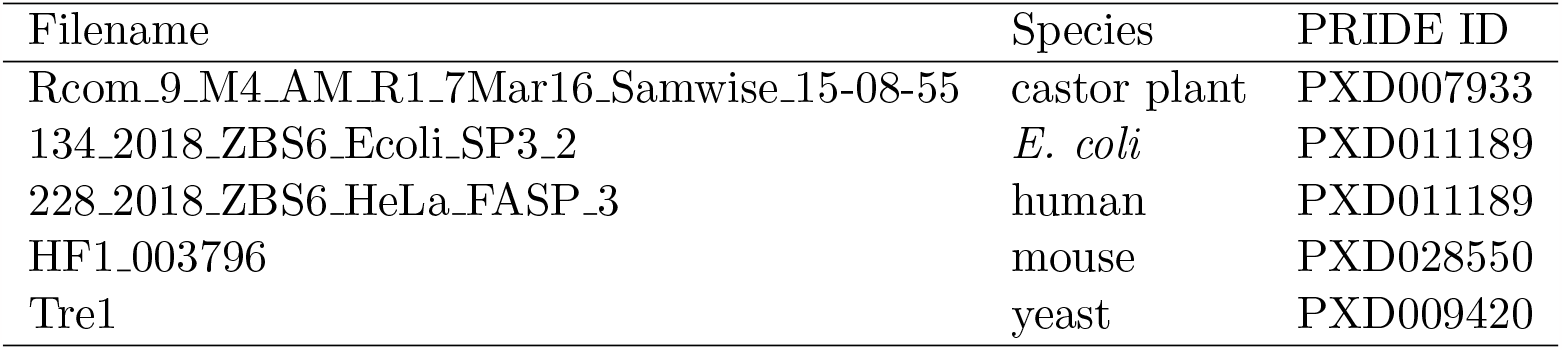
Datasets used in this manuscript.

We searched each run against its respective proteome using eight different score functions from five different database search engines. The following search engines were used: Tide version 4.1,^24^ MSGF+ version v2021.09.06,^17^ MSFragger version 3.5,^27^ MSAmanda version 2.0.0.18350,^28^ and Comet version 2022.01 rev. 1 (11cb28f).^25,26,34^ Searches using Tide employed four different score functions: XCorr,^24^ XCorr p-value,^35^ combined p-value,^36^ and the Tailor score.^37^ For the searches with Tide, we used the default parameters except “concat=F” and “peptide-list=T”. With MSGF+, all parameters were set to their default values except for “inst 3” and “tda 1”. We used all default parameters for MSAmanda and MSFragger, and the concatenated target-decoy database used in MSFragger was generated by Philosopher.^38^ For Comet we used the default parameters found in comet.params.high-high except that “decoy_search=2”, “output_txtfile=1”, “output_mzidentmlfile”, and “num_output_lines=1”. The output of these database searches were provided to Crema for FDR estimation.

In addition to the database searches above, we also performed additional Tide searches to compare the performance between an explicit and implicit target-decoy pairing. We used the same five runs and four score functions that were previously used. In addition, the parameters that were used remained exactly the same except that we allowed for up to five methionine oxidations. The explicit target-decoy pairing was provided by the tide-index tool.

## 3 Results

### 3.1 Tutorial: Computing FDR estimates at the PSM, peptide and protein levels

In addition to standardizing how TDC estimates the FDR, we intend Crema to be a straightforward program to install and use. To that end, we give users the ability to run Crema through a command-line interface or via a Python API. In addition, we provide here a tutorial of how to run Crema, and additional usage information can be found at https://crema-ms.readthedocs.io.

Estimating the FDR among a set of PSMs using the Crema Python API is simple and can be performed using only a couple lines of code. We provide example code showing how to estimate the FDR on database search results from Tide (Figure 1). After importing the Crema package, the PSMs are read using the “read_tide” function. This function is given a list of search results. We note that these input files can be either system paths or Pandas dataframes. In the provided example, `target_file` contains target PSMs and `decoy_file` contains decoy PSMs (Figure 1). In addition, if a file that explicitly pairs target and decoy sequences exist, it can be passed by the “pairing_file_name” argument. Finally, the “read_tide” function can be switched out for other functions, such as “read_msgf” and “read_msfragger”, depending on which database search engine was used.

**Figure 1.**
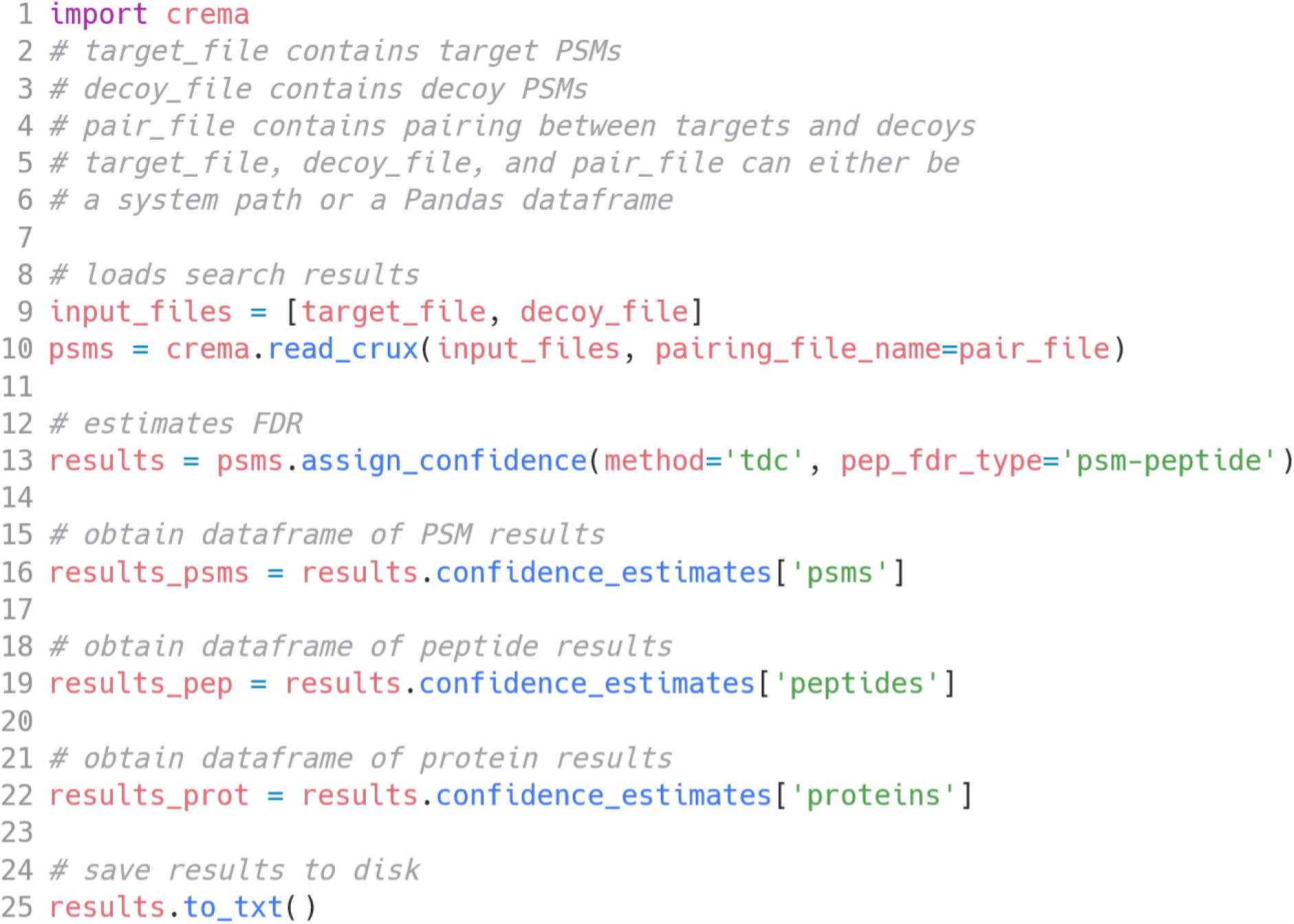
Code for running Crema. Example code for computing FDR estimates at the PSM, peptide, and protein levels using Crema on output from Tide.

After reading and storing the PSMs, the FDR is estimated using the “assign_confidence” function. The user has three choices, via the “pep_fdr_type” argument, for estimating peptide level FDR: “psm-only”, “peptide-only”, and “psm-peptide”. The “assign_confidence” returns a Confidence object that contains the list of discoveries at a given FDR threshold at the psm, peptide, and protein level. The discoveries at a single level can be obtained as a Pandas dataframe using the “confidence_estimates” function. Alternatively, results can be returned as a text files (one each for PSM, peptide and protein level FDR) using the “to txt” function. In addition, instead of discoveries, Crema can also provide q-values. As noted, we strongly discourage the use of q-values for anything other than benchmarking.

### 3.2 Comparison of peptide-level FDR control procedures

We first compared the three peptide-level FDR control procedures implemented within Crema. Note that, due to the inability to match targets to decoys with some search engines, we report results for MSGF+, MSFragger, and MSAmanda using only the psm-only method.

Our results provide evidence, as previously shown,^5^ that psm-peptide outperforms both peptide-only and psm-only, and that peptide-only outperforms psm-only (Figure 2). This trend held across all runs and score functions over the entire FDR range of 0–10%. The sole exception was for runs analyzed by Comet. For these searches, peptide-only and psm-only had very similar performance (far left column of Figure 2). At a 1% FDR, we found that psm-peptide outperformed psm-only and peptide-only by an average of 17.14% and 9.52% peptide detections, respectively. Note that this average does not include data from the castor run analyzed by XCorr, because psm-only did not obtain any confident detections at 1% FDR.

**Figure 2.**
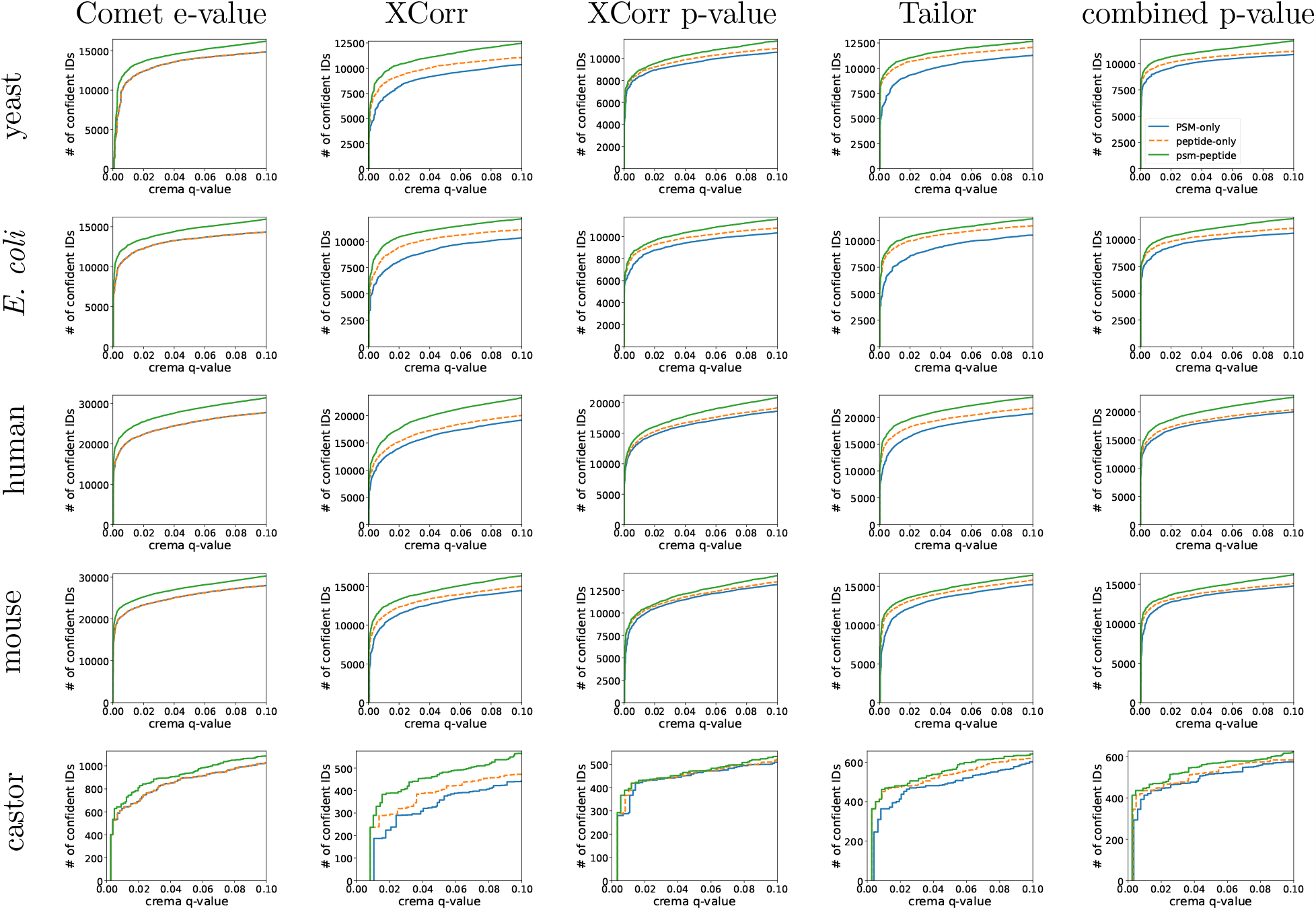
Peptide-level Crema output. Each panel shows the number of peptide detections as a function of FDR threshold after using Crema. Each row of panels represents a run from a different species and each column represents a different score function.

As we have noted, the only peptide-level FDR control procedure that is compatible with the decoys produced by MSGF+, MSFragger, or MSAmanda is psm-only. Accordingly, we show the performance curves for these search engines in Figure 3. Importantly, we note that, in these results, one cannot easily compare the number of discoveries from different search engines against each other. This is because different search engines use different decoy generation methods. We also did not verify that all of the user-specified parameters were exactly comparable across search engines. Thus, any observed difference in performance may be a result of a combination of factors.

**Figure 3.**
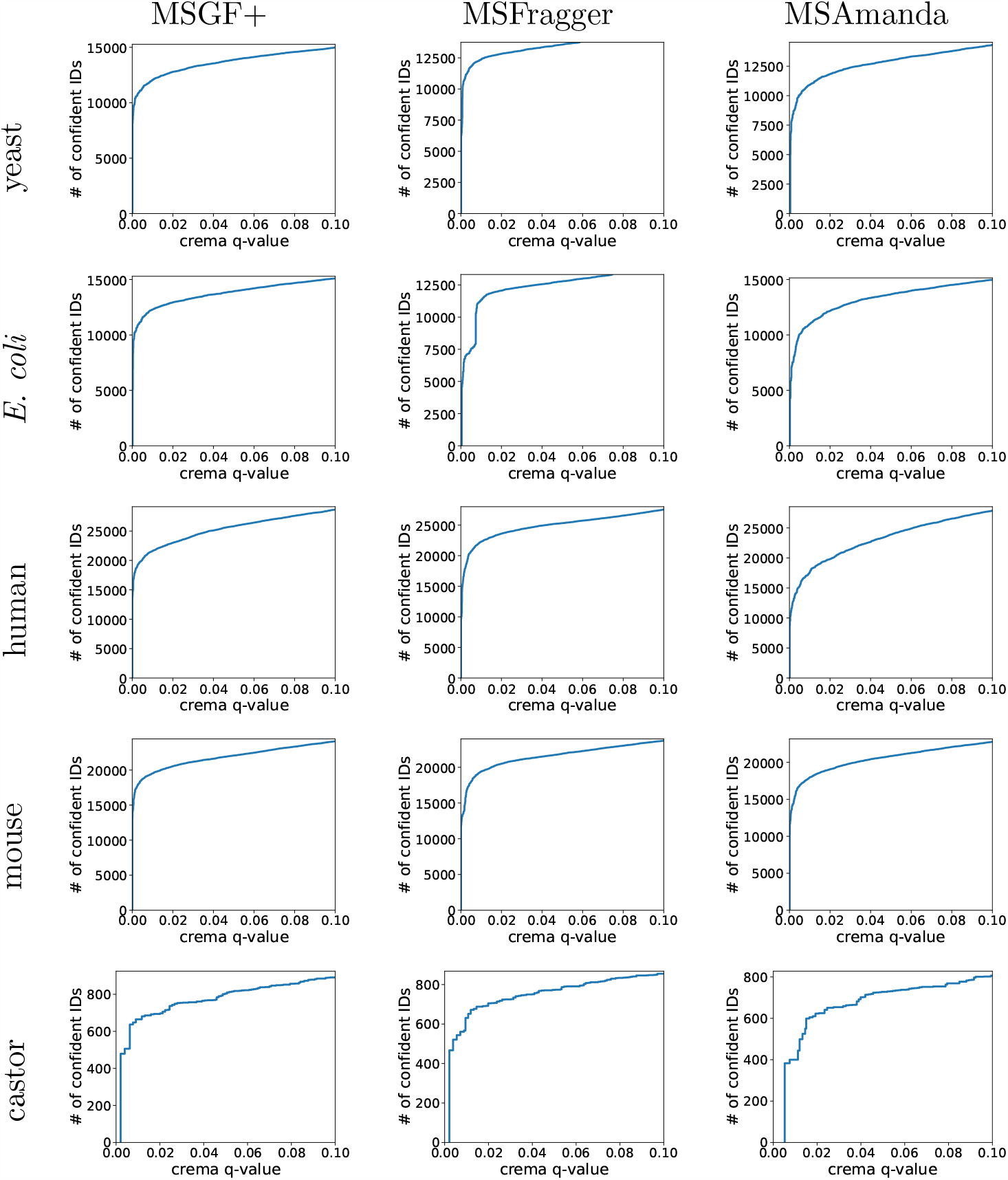
Peptide-level Crema output for MSGF+, MSFragger and MSAmanda. Each panel shows the number of peptide detections, using “psm-only”, as a function of FDR threshold estimated by Crema. Each row of panels represents a run from a different species, and each column represents a different database search engine.

The best-performing peptide-level FDR estimation procedure, psm-peptide, requires a matching between a target peptide and its corresponding decoy peptide. With Tide, this matching can be provided by the user or can be inferred automatically by Crema. We verified that both approaches yield similar results, for a variety of Tide score functions (Supplementary Figure S3).

Finally, within Crema, there is one method estimating the FDR at the PSM level and one method for the protein level. We systematically examined the performance of these two methods for all score functions and runs (Supplementary Figures S1–S5). Note that we also considered a variant of the protein-level FDR estimation procedure, in which a protein score is the sum (rather than the maximum) of the unique peptides scores. Ultimately, we chose not to implement this method within Crema because it performed significantly worse than the original method (Supplementary Figure S4–S5).

## 4 Discussion

In this work, we introduce an open-source Python tool, Crema, that implements several TDC-based methods for estimating the FDR. Crema can currently be used with five search engines, and it can provide an easy way to obtain accurate and robust FDR estimates.

The results in Figure 2 support the idea that performing “double competition” between targets and decoys, as in the psm-peptide procedure, provides a substantial boost in statistical power to detect peptides. This is an important but not novel observation, as we have previously reported similar results.^5^ Furthermore, it is worth noting that our previous paper includes additional empirical evidence (in the form of entrapment experiments^39^) that psm-peptide’s increased power is not due to lax control of the FDR.

Several decoy-free procedures for estimating the FDR have been proposed,^40,41^ but these are not implemented within Crema. We focused on TDC-based methods because TDC is the most commonly used method for estimating the FDR.

Although Crema can help to ensure that practitioners employ valid FDR control procedures, it does not solve all possible analytical issues. For example, if a given dataset is searched against a problematic decoy database, then Crema cannot help. In addition, Crema is unable to help with incorrect post-processing or biological interpretation. Finally, Crema does not employ machine learning approaches for estimating the FDR employed by methods such as Percolator,^13^ Mokapot,^42^ and PeptideProphet.^22^

In the future, we hope to improve Crema further. We plan, for example, to add a graphical user interface to the tool. We also plan to expand support for additional search engines and output formats. Users with specific feature requests are invited to submit them via the Crema issue tracker on Github.

## Acknowledgments

Some of the research described in this paper was conducted under the Laboratory Directed Research and Development Program at Pacific Northwest National Laboratory, a multiprogram national laboratory operated by Battelle for the U.S. Department of Energy. Andy Lin is grateful for the support of the Linus Pauling Distinguished Postdoctoral Fellowship program. Pacific Northwest National Laboratory is a multiprogram national laboratory operated by Battelle Memorial Institute for the United States Department of Energy under contract DE-AC06-76RLO.

## Data Availability

No new data were created or analyzed for this study.

## Supporting information

- **Supplemental File S1:** PDF containing Supplemental Figures and Tables.

## Footnotes

1 Terms in boldface are defined in Box 1.

## References

[1] J. E. Elias and S. P. Gygi. Target-decoy search strategy for increased confidence in large-scale protein identifications by mass spectrometry. Nature Methods, 4(3):207–214, 2007.

[2] K. He, Y. Fu, W.-F. Zeng, L. Luo, H. Chi, C. Liu, L.-Y. Qing, R.-X. Sun, and S.-M. He. A theoretical foundation of the target-decoy search strategy for false discovery rate control in proteomics. arXiv, 2015. https://arxiv.org/abs/1501.00537.

[3] K. Jeong, S. Kim, and N. Bandeira. False discovery rates in spectral identification. BMC Bioinformatics, 13(Suppl. 16):S2, 2012.

[4] M. Wilhelm, J. Schlegl, H. Hahne, A. Moghaddas Gholami, M. Lieberenz, M. M Savitski, E. Ziegler, L. Butzmann, S. Gessulat, H. Marx, T. Mathieson, S. Lemeer, K. Schnatbaum, U. Reimer, H. Wenschuh, M. Mollenhauer, J. Slotta-Huspenina, J. Boese, M. Bantscheff, A. Gerstmair, F. Faerber, and B. Kuster. Mass-spectrometry-based draft of the human proteome. Nature, 509(7502):582–587, 2014.

[5] A. Lin, T. Short, W. S. Noble, and U. Keich. Improving peptide-level mass spectrometry analysis via double competition. Journal of Proteome Research, 21(10):2412–2420, 2022.

[6] R. F. Barber and Emmanuel J. Candès. Controlling the false discovery rate via knockoffs. The Annals of Statistics, 43(5):2055–2085, 2015.

[7] J. E. Elias and S. P. Gygi. Target-decoy search strategy for mass spectrometry-based proteomics. Methods in Molecular Biology, 604(55–71), 2010.

[8] G. Wang, W. W. Wu, Z. Zhang, S. Masilamani, and R.-F. Shen. Decoy methods for assessing false positives and false discovery rates in shotgun proteomics. Analytical Chemistry, 81(1):146–159, 2009.

[9] W. H. Vensel, F. M. DuPont, S. Sloane, and S. B. Altenbach. Effect of cleavage enzyme, search algorithm and decoy database on mass spectrometric identification of wheat gluten proteins. Phytochemistry, 72(10):1154–1161, 2011.

[10] J. M. Moosa, S. Guan, M. F. Moran, and B. Ma. Repeat-preserving decoy database for false discovery rate estimation in peptide identification. Journal of Proteome Research, 19(3):1029–1036, 2020.

[11] L. I. Levitsky, M V. Ivanov, A. A. Lobas, and M. V. Gorshkov. Unbiased false discovery rate estimation for shotgun proteomics based on the target-decoy approach. Journal of Proteome Research, 16(2):393–397, 2017.

[12] A. I. Nesvizhskii, A. Keller, E. Kolker, and R. Aebersold. A statistical model for identifying proteins by tandem mass spectrometry. Analytical Chemistry, 75:4646–4658, 2003.

[13] L. Käll, J. Canterbury, J. Weston, W. S. Noble, and M. J. MacCoss. A semi-supervised machine learning technique for peptide identification from shotgun proteomics datasets. Nature Methods, 4:923–25, 2007.

[14] J. Freestone, T. Short, W. S. Noble, and U. Keich. Group-walk: a rigorous approach to group-wise false discovery rate analysis by target-decoy competition. Bioinformatics, 38(Supplement 2):ii82–ii88, 09 2022.

[15] W. S. Noble. Mass spectrometrists should only search for peptides they care about. Nature Methods, 12(7):605–608, 2015.

[16] A. Sticker, L. Martens, and L. Clement. Mass spectrometrists should search for all peptides, but assess only the ones they care about. Nature Methods, 14(7):643–644, 2017.

[17] S. Kim and P. A. Pevzner. MS-GF+ makes progress toward a universal database search tool for proteomics. Nature Communications, 5:5277, 2014.

[18] C. Y. Park, A. A. Klammer, L. Käll, M. P. MacCoss, and W. S. Noble. Rapid and accurate peptide identification from tandem mass spectra. Journal of Proteome Research, 7(7):3022–3027, 2008.

[19] A. A. Goloborodko, L. I. Levitsky, M. V. Ivanov, and M. V. Gorshkov. Pyteomics–a Python framework for exploratory data analysis and rapid software prototyping in proteomics. Journal of the American Society for Mass Spectrometry, 24(2):301–304, Feb 2013.

[20] L. I. Levitsky, J. A. Klein, M. V. Ivanov, and M. V. Gorshkov. Pyteomics 4.0: Five Years of Development of a Python Proteomics Framework. Journal of Proteome Research, 18(2):709–714, Feb 2019.

[21] H. L. Röst, T. Sachsenberg, S. Aiche, C. Bielow, H. Weisser, F. Aicheler, S. Andreotti, H. Ehrlich, P. Gutenbrunner, E. Kenar, X. Liang, S. Nahnsen, L. Nilse, J. Pfeuffer, G. Rosenberger, M. Rurik, U. Schmitt, J. Veit, M. Walzer, D. Wojnar, W. E. Wolski, O. Schilling, J. S. Choudhary, L. Malmström, R. Aebersold, K. Reinert, and O. Kohlbache. OpenMS: a flexible open-source software platform for mass spectrometry data analysis. Nature Methods, 13(9):741, August 2016.

[22] A. Keller, A. I. Nesvizhskii, E. Kolker, and R. Aebersold. Empirical statistical model to estimate the accuracy of peptide identification made by MS/MS and database search. Analytical Chemistry, 74:5383–5392, 2002.

[23] E. Debrie, M. Malfait, R. Gabriels, A. Declerq, A. Sticker, L. Martens, and L. Clement. Quality Control for the Target Decoy Approach for Peptide Identification. J Proteome Res, 22(2):350–358, Feb 2023.

[24] B. Diament and W. S. Noble. Faster SEQUEST searching for peptide identification from tandem mass spectra. Journal of Proteome Research, 10(9):3871–3879, 2011.

[25] J. K. Eng, T. A. Jahan, and M. R. Hoopmann. Comet: an open source tandem mass spectrometry sequence database search tool. Proteomics, 13(1):22–24, 2012.

[26] J. K. Eng, M. R. Hoopmann, T. A. Jahan, J. D. Egertson, W. S. Noble, and M. J. MacCoss. A deeper look into Comet–implementation and features. Journal of the American Society for Mass Spectrometry, 26(11):1865–1874, Nov 2015.

[27] A. T. Kong, F. V. Leprevost, D. M. Avtonomov, D. Mellacheruvu, and A. I. Nesvizhskii. MSFragger: ultrafast and comprehensive peptide identification in mass spectrometry-based proteomics. Nature Methods, 14(5):513–520, 2017.

[28] V. Dorfer, P. Pichler, T. Stranzl, J. Stadlmann, T. Taus, S. Winkler, and K. Mechtler. MSAmanda, a universal identification algorithm optimized for high accuracy tandem mass spectra. Journal of Proteome Research, 13(8):3679–3684, 2014.

[29] Andrew Rajchert and Uri Keich. Controlling the false discovery rate via competition: Is the +1 needed? Statistics & Probability Letters, 197:109819, 2023.

[30] Y. Perez-Riverol, A. Csordas, J. Bai, M. Bernal-Llinares, S. Hewapathirana, D. J. Kundu, A. Inuganti, J. Griss, G. Mayer, M. Eisenacher, E. Pérez, J. Uszkoreit, J. Pfeuffer, T. Sachsenberg, S. Yilmaz, S. Tiwary, J. Cox, E. Audain, M. Walzer, A. F. Jarnuczak, T. Ternent Brazma, and J. A. Vizcaíno. The PRIDE database and related tools and resources in 2019: improving support for quantification data. Nucleic Acids Res, 47(D1):D442–D450, Jan 2019.

[31] M. C. Chambers, B. Maclean, R. Burke, D. Amodei, D. L. Ruderman, S. Neumann, L. Gatto Fischer, B. Pratt, J. Egertson, K. Hoff, D. Kessner, N. Tasman, N. Shulman, B. Frewen, T. A. Baker, M. Y. Brusniak, C. Paulse, D. Creasy, L. Flashner, K. Kani, C. Moulding, S. L. Seymour, L. M. Nuwaysir, B. Lefebvre, F. Kuhlmann, J. Roark, P. Rainer, S. Detlev, T. Hemenway, A. Huhmer, J. Langridge, B. Connolly, T. Chadick, K. Holly, J. Eckels, E. W. Deutsch, R. L. Moritz, J. E. Katz, D. B. Agus, M. J. MacCoss, D. L. Tabb, and P. Mallick. A cross-platform toolkit for mass spectrometry and proteomics. Nature Biotechnology, 30(10):918–920, 2012.

[32] UniProt Consortium. UniProt: a hub for protein information. Nucleic Acids Research, page gku989, 2014.

[33] UniProt Consortium. UniProt: a worldwide hub for protein knowledge. Nucleic Acids Research, pages D506–D515, 2019.

[34] J. K. Eng, B. Fischer, J. Grossman, and M. J. MacCoss. A fast SEQUEST cross correlation algorithm. Journal of Proteome Research, 7(10):4598–4602, 2008.

[35] J. J. Howbert and W. S. Noble. Computing exact p-values for a cross-correlation shotgun proteomics score function. Molecular and Cellular Proteomics, 13(9):2467–2479, 2014.

[36] A. Lin, J. J. Howbert, and W. S. Noble. Combining high-resolution and exact calibration to boost statistical power: A well-calibrated score function for high-resolution MS2 data. Journal of Proteome Research, 17:3644–3656, 2018.

[37] P. Sulimov and A. Kertész-Farkas. Tailor: A nonparametric and rapid score calibration method for database search-based peptide identification in shotgun proteomics. Journal of Proteome Research, 19(4):1481–1490, 2020.

[38] F. da Veiga Leprevost, S. E. Haynes, D. M. Avtonomov, H. Y. Chang, A. K. Shanmugam, D. Mellacheruvu, A. T. Kong, and A. I. Nesvizhskii. Philosopher: a versatile toolkit for shotgun proteomics data analysis. Nature Methods, 17(9):869–870, Sep 2020.

[39] V. Granholm, J. F. Navarro, W. S. Noble, and L. Käll. Determining the calibration of confidence estimation procedures for unique peptides in shotgun proteomics. Journal of Proteomics, 80(27):123–131, 2013.

[40] Y. Couté, C. Bruley, and T. Burger. Beyond target-decoy competition: Stable validation of peptide and protein identifications in mass spectrometry-based discovery proteomics. Analytical Chemistry, 92(22):14898–14906, 2020.

[41] Y. Peng, S. Jain, Y. F. Li, M. Greguš, A. R. Ivanov, O. Vitek, and P. Radivojac. New mixture models for decoy-free false discovery rate estimation in mass spectrometry proteomics. Bioinformatics, 36(Supplement 2):i745–i753, 2020.

[42] W. E. Fondrie and W. S. Noble. mokapot: Fast and flexible semi-supervised learning for peptide detection. Journal of Proteome Research, 20(4):1966–1971, 2021.

