## Supplemental File for "Target-decoy false discovery rate estimation using Crema"

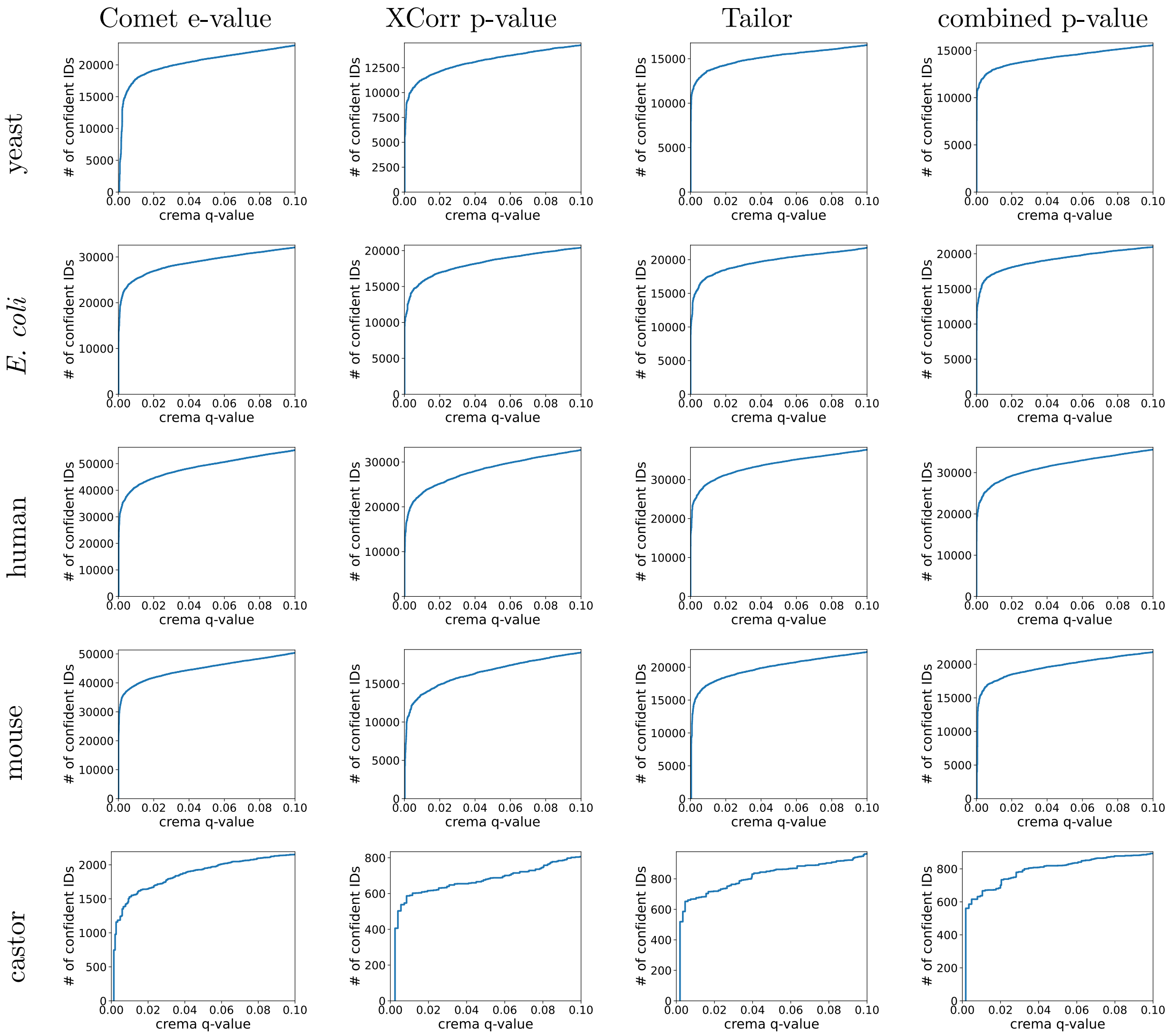


Figure S1: **PSM-level Crema output using Tide.** Each panel shows the number of PSM detections as a function of FDR threshold, estimated using Crema. Each row of panels represents a run from a different species, and each column represents a different score function available from Tide.

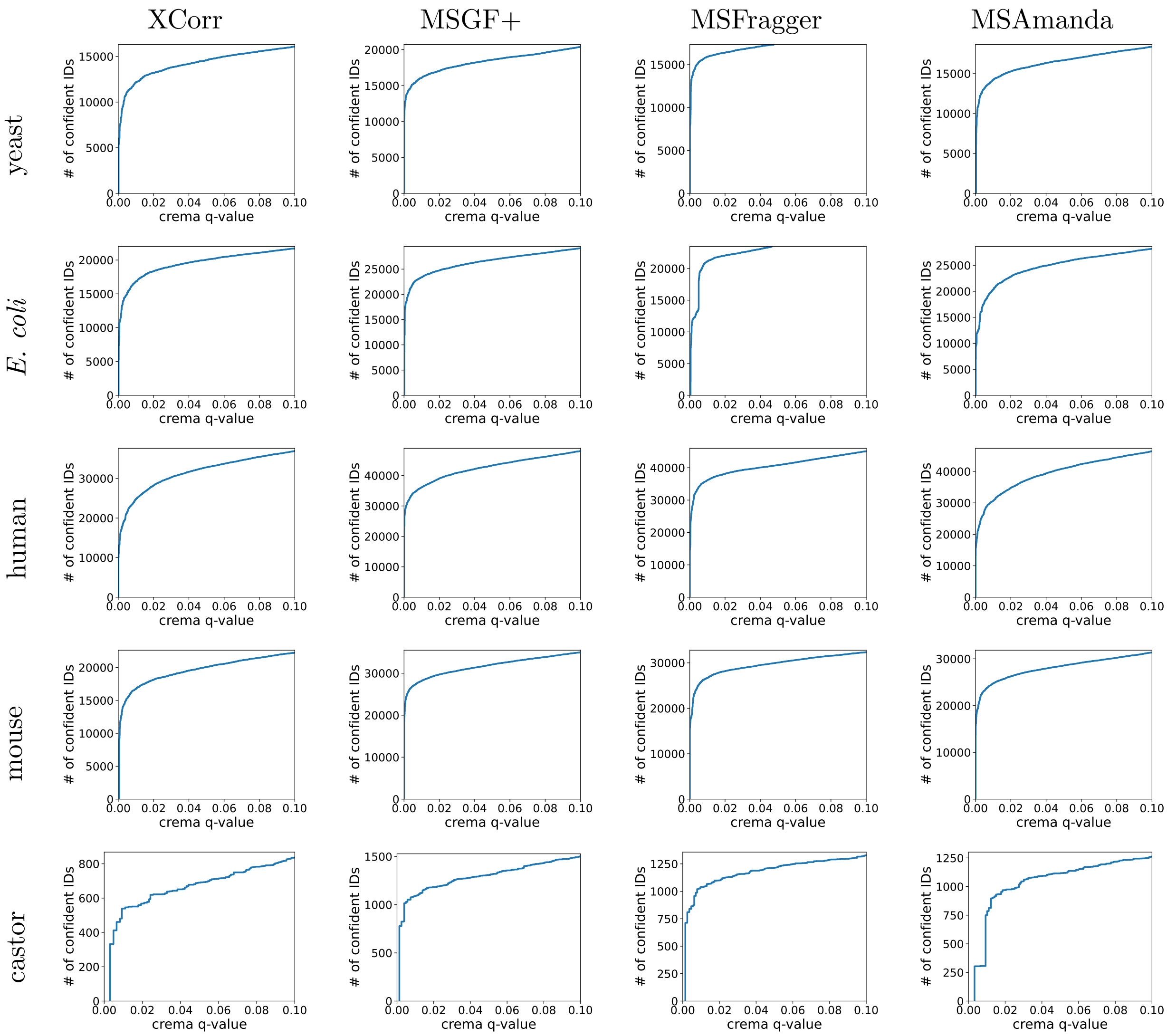


Figure S2: **PSM-level Crema output using other search engines.** Each panel shows the number of accepted PSMs as a function of FDR threshold estimated via TDC by Crema. Each row of panels represents a run from a different species, and each column represents a different database search engine.

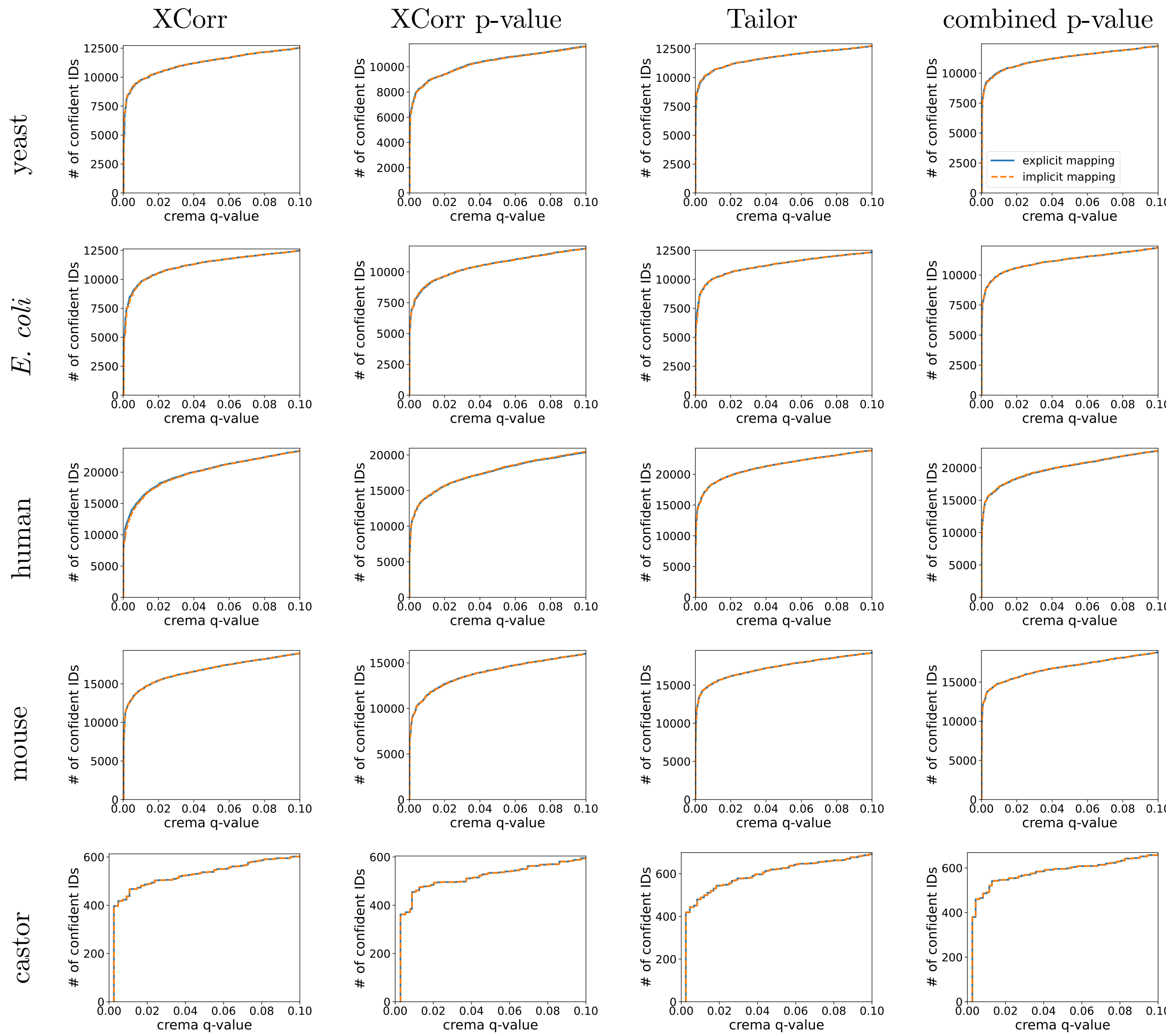


Figure S3: **Comparison between explicit and implicit mapping for peptide-level FDR.** Each panel shows the number of peptide detections, after using the psm-peptide method, as a function of FDR threshold estimated by Crema with explicit or implicit target-decoy pairing. Each row of panels represents a run from a different species, and each column represents a different score function available from Tide.

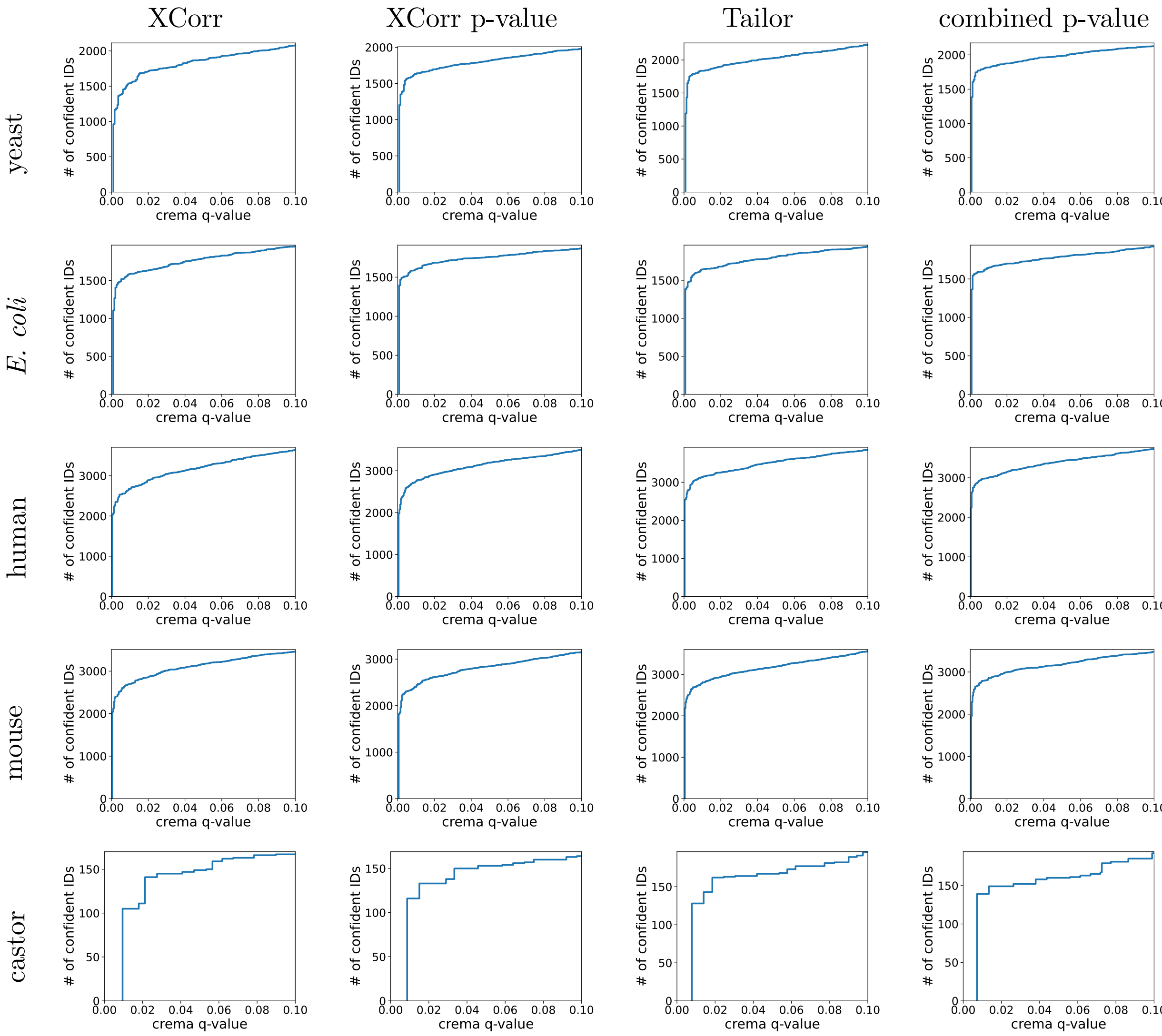


Figure S4: **Protein-level Crema output using Tide.** Each panel shows the number of protein detections as a function of FDR threshold estimated by Crema. Each row of panels represents a run from a different species, and each column represents a different score function available from Tide.

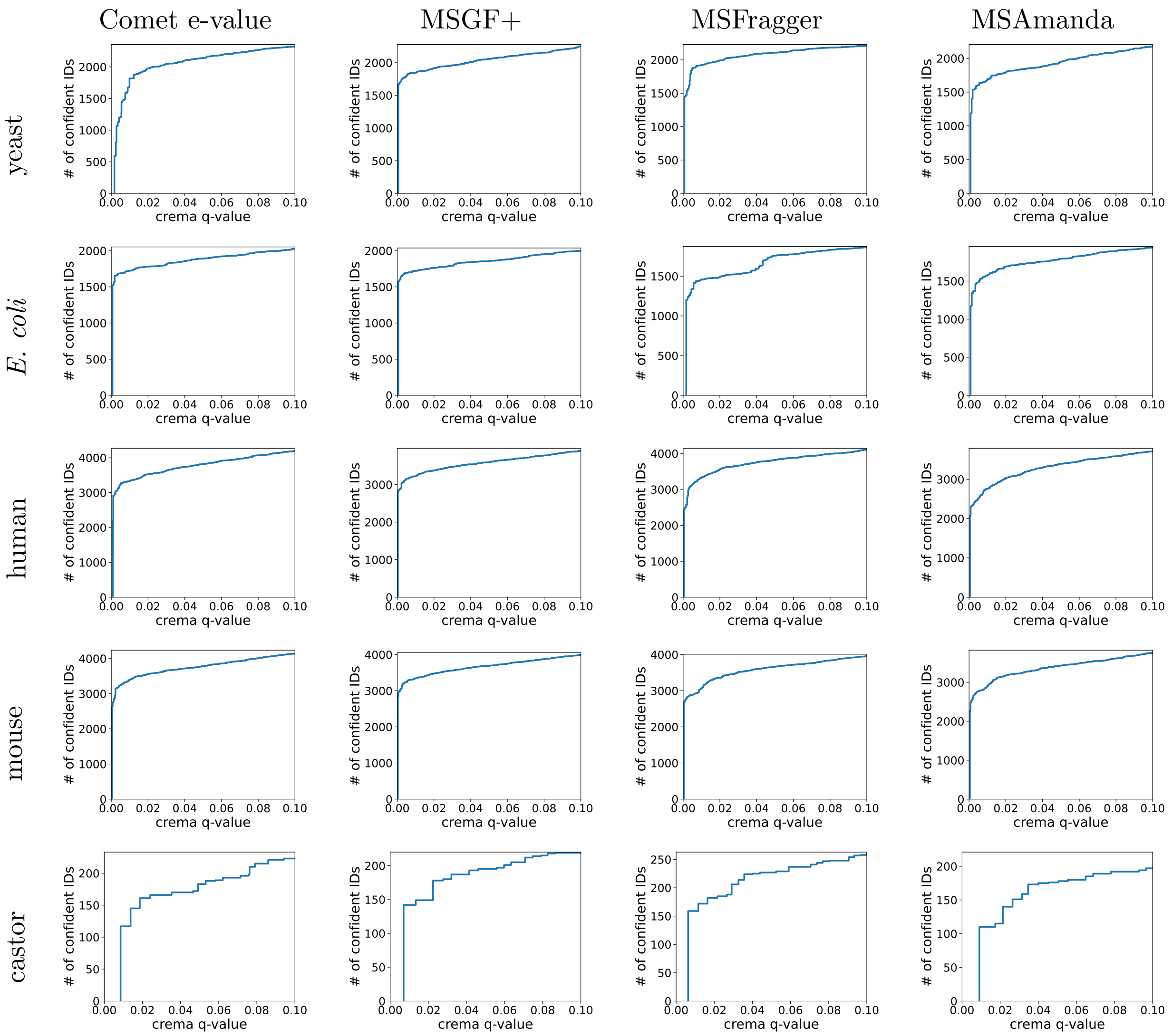


Figure S5: **Protein-level Crema output using other search engines.** Each panel shows the number of protein detections as a function of FDR threshold estimated by Crema. Each row of panels represents a run from a different species, and each column represents a different database search engine.

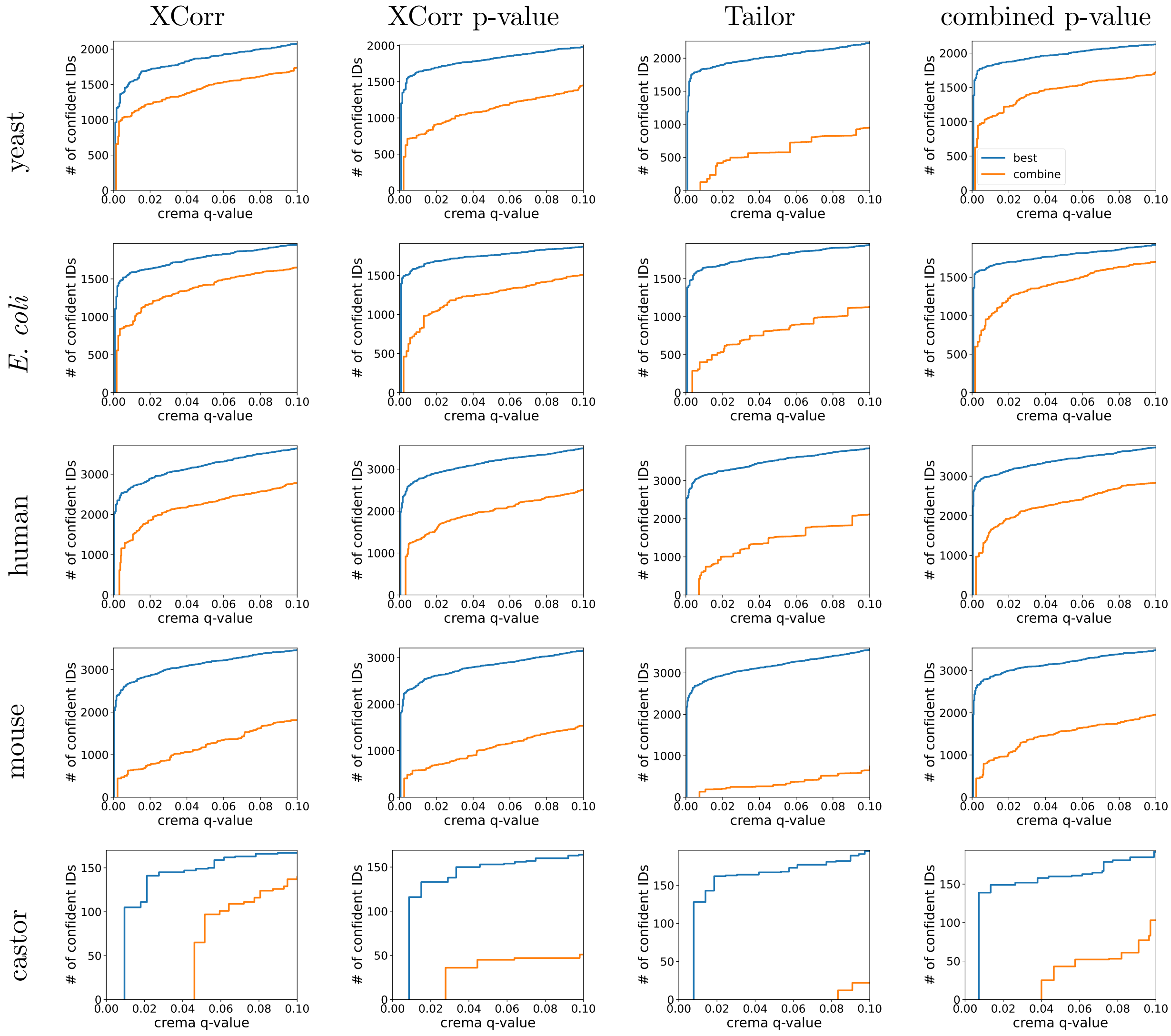


Figure S6: **Comparison of protein-level FDR methods using Tide.** Each panel shows the number of protein detections as a function of FDR threshold for two protein-level FDR estimation methods: "best" assigns each protein the maximum associated PSM score, whereas "combined" assigns the sum of the scores associated with unique peptides. Each row of panels represents a run from a different species, and each column represents a different score function available from Tide.

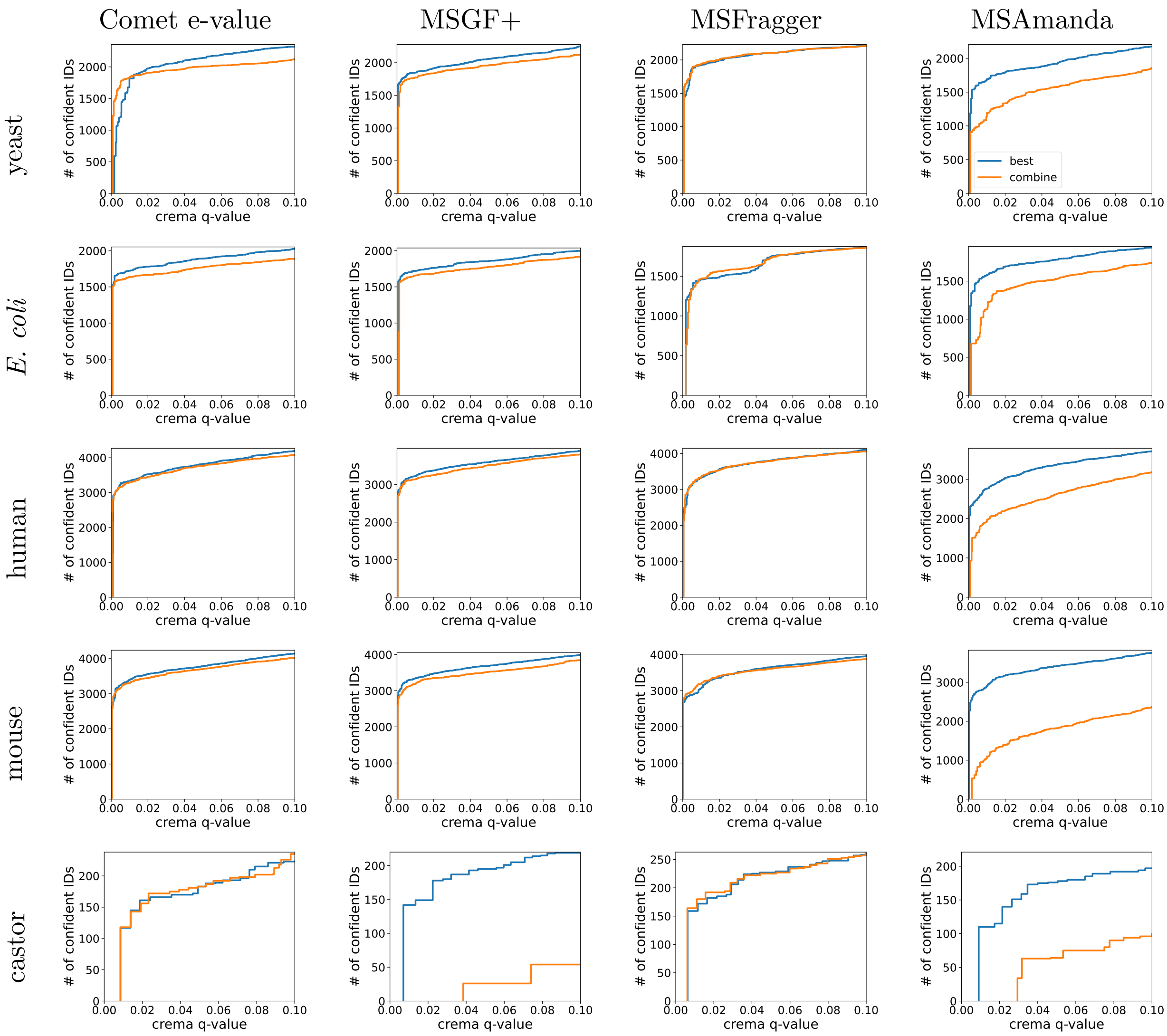

Figure S7: **Comparison of protein-level FDR methods using various search engines.** Each panel shows the number of protein detections as a function of FDR threshold for two protein-level FDR estimation methods. Each row of panels represents a run from a different species, and each column represents a different database search engine.
